# Early Pulmonary Fibrosis is Defined by Niche- and Cell-Specific Molecular Programs

**DOI:** 10.64898/2026.05.28.727955

**Authors:** Alan Waich, Scott A. Ochsner, Julian A. Villalba, Jonathan A. Rose, Juan D. Cala Garcia, Juan D. Zuluaga, Neil J. Mckenna, Maria E. Ruiz Echartea, Chao He, Lindsay J. Celada, Konstantin Tsoyi, Laura F. Gonzalez-Cuevas, Andrea Galecio Chao, Aurelien Justet, Stefan W. Ryter, Wendy J. Introne, Naftali Kaminski, David A. Schwartz, Benjamin A. Raby, Gary M. Hunninghake, Bernadette R. Gochuico, Cristian Coarfa, Ivan O. Rosas

## Abstract

**Rationale:** Preclinical familial pulmonary fibrosis (FPF) represents an early stage of fibrotic lung disease, yet the compartment- and cell-specific molecular programs preceding fibrosis remain poorly understood.

**Objective:** To define spatially organized molecular signatures associated with preclinical FPF and identify tissue-informed circulating biomarkers linked to early fibrotic remodeling.

**Methods:** We performed integrated multi-omic profiling of histologically preserved and remodeled lung regions from subjects with preclinical FPF, Idiopathic Pulmonary Fibrosis (IPF), and controls using spatial transcriptomics, single-nucleus RNA sequencing (snRNAseq), and blood proteomics. Differential expression and pathway enrichment analyses were performed across spatial compartments and epithelial cell states.

**Results:** Histologically preserved lung regions in preclinical FPF demonstrated transcriptional abnormalities including stress-response, ciliary, and extracellular matrix-associated programs despite minimal architectural distortion. Spatial analyses identified alterations in alveolar niche molecular programs accompanied by increasing profibrotic signaling across preserved and tissue remodeled lung compartments. Compared with advanced IPF, preclinical FPF retained epithelial repair and surfactant-associated signatures. Integration with snRNAseq demonstrated enrichment of alveolar and airway epithelial cell dysregulated states associated with transitional phenotypes previously implicated in IPF. Compartment- and epithelial-associated transcriptional signatures identified in lung tissue were partially represented in the peripheral blood.

**Conclusion:** Preclinical FPF is characterized by compartment- and cell-specific molecular programs that precede established fibrosis. We identified distinct alveolar, airway, and vascular molecular signatures and epithelial remodeling states represented in the peripheral blood. These findings provide an initial framework for molecular classification of early stages of pulmonary fibrosis and support future studies evaluating minimally invasive approaches for disease stratification and precision therapeutics.

**At a Glance Commentary:** *Scientific Knowledge on the Subject:* The molecular events preceding a diagnosis of pulmonary fibrosis remain poorly understood. Most mechanistic studies in Idiopathic Pulmonary Fibrosis (IPF) have relied on end-stage explanted lungs, limiting insight into the compartment- and cell-specific molecular programs associated with early stages of pulmonary fibrosis.

*What this study adds to the field:* Using integrated spatial transcriptomics, single-cell sequencing, and peripheral blood proteomic profiling, we demonstrate that preclinical familial pulmonary fibrosis (FPF) is characterized by compartment- and cell-specific molecular programs that precede clinically detectable fibrosis. Spatial analyses identified distinct alveolar, airway, and vascular molecular signatures, while single cell analysis confirmed the presence of epithelial dysregulated states. These signatures are partially represented in the peripheral blood. Our findings provide an initial framework for biologically informed classification of early stages of pulmonary fibrosis and future minimally invasive approaches for disease stratification.

## INTRODUCTION

Idiopathic Pulmonary Fibrosis (IPF) is a progressive fibrotic lung disease characterized by spatially heterogeneous remodeling of the lung parenchyma, including fibroblastic foci, dense fibrotic regions, and honeycombing, amid areas of relatively histologically preserved architecture. Similar histologic features occur in early stages of familial pulmonary fibrosis (FPF), suggesting that shared molecular mechanisms are preserved across fibrotic diseases^1^. However, mechanistic insights in pulmonary fibrosis research are largely derived from explanted lungs^2^, in which extensive tissue remodeling may obscure the molecular programs that initiate fibrosis and precede irreversible architectural destruction. Consequently, the earliest biological events driving fibrotic lung disease remain unresolved.

Recent advances in computed tomography (CT)-based screening of individuals at risk for pulmonary fibrosis, particularly first-degree relatives of patients with FPF, have improved the detection of interstitial lung abnormalities (ILAs)^3–5^. Radiographic abnormalities identified in asymptomatic individuals share genetic and biologic overlap with established IPF, including associations with MUC5B and other fibrosis susceptibility loci^6,7^. Longitudinal studies reveal temporal progression of fibrotic ILAs, supporting that IPF begins years before clinical diagnosis^3^. However, ILAs are mostly defined by imaging phenotypes which fail to capture the heterogeneity of early fibrotic remodeling^8–10^. Consequently, the compartment- and cell-specific molecular programs that determine preclinical fibrosis may identify molecular programs that could be targeted to prevent the progression of pulmonary fibrosis.

Identification of peripheral blood biomarkers associated with pulmonary fibrosis have improved the identification of early disease^11,12^. Circulating protein signatures have been associated with established IPF^11,13^, and recently, with the presence and progression of ILAs^14^. However, these biomarkers may incompletely capture biological heterogeneity of preclinical pulmonary fibrosis. Furthermore, similar radiographic phenotypes may arise from distinct epithelial, stromal, vascular, or immune molecular programs.

Recent advances in single-cell and spatial transcriptomic profiling have defined the cellular architecture of pulmonary fibrosis^15–17^. Fibrotic remodeling is characterized by spatially organized niches involving epithelial, stromal, endothelial, and immune cell populations^17–19^. Transitional epithelial states, including alveolar intermediate and aberrant basaloid populations, have been associated with extracellular matrix (ECM) remodeling and failed epithelial repair in IPF^16,20^. Spatial transcriptomic analyses of explanted lungs at end-stage disease have identified distinct molecular programs localized to specific histologic niches^18,21^.

To investigate whether compartment- and cell-specific molecular programs precede established fibrotic remodeling in IPF, we performed multi-omic studies in preclinical FPF subjects. Using integrated spatial transcriptomic and single-nuclei approaches, we interrogated histologically preserved and remodeled lung regions to define compartment- and cell-specific molecular signatures associated with early fibrotic remodeling; and investigated whether these signatures are represented in the peripheral blood. This study will enable molecular classification of early-stage pulmonary fibrosis and support future development of minimally invasive approaches for preventive and precision medicine.

## MATERIALS AND METHODS

Additional detail on all methods is provided in the **Supplementary Methods**.

### Lung tissue samples

Lung tissue was obtained from open lung biopsies of eight individuals (four female, four male, average age = 55.35 years) with preclinical FPF, defined by radiographic evidence of ILA and absence of clinical symptoms. These were collected as part of a larger cohort of kindreds of patients affected with pulmonary fibrosis^1^ enrolled in protocols 99-H-0068, 04-HG-0211 and/or NCT00084305 approved by the National Heart, Lung, and Blood Institute (NHLBI) and/or the National Human Genome Research Institute (NHGRI) respective Institutional Review Boards (IRB). Additionally, lung explants were collected from four individuals with end-stage IPF undergoing lung transplantation. The diagnosis of IPF was established according to the American Thoracic Society/European Respiratory Society (ATS/ERS) guidelines^22^. Age-matched control samples were obtained from organ donors whose lungs were rejected for transplantation. Lung sample collection was approved by the Baylor College of Medicine (BCM) IRB (H-46823). Clinical and demographic characteristics of study subjects are listed in **Table S1.**

### Bruker Spatial Biology GeoMx Digital Spatial Profiler

Spatial transcriptomic profiling was performed using a Bruker Spatial Biology GeoMx Digital Spatial Profiler (DSP). RNA slides were prepared using the manufacturer’s protocols. 5 μm tissue sections were hybridized with the GeoMx Whole Transcriptome Atlas probe set and stained with fluorescent antibodies (pan-cytokeratin, SYTO13 and CD45). Regions of Interest (ROI) were selected by a board-certified and fellowship-trained pulmonary pathologist (J.A.V.) guided by hematoxylin-eosin (H&E) and fluorescent antibody staining. Four to five regions were selected per ROI type per sample to capture sufficient spatial heterogeneity and produce stable variance estimates (**Table S2**). Serial ultraviolet illumination of each region was used to collect the probe barcodes into a 96-well collection plate. Each ROI was then indexed using an Illumina dual indexing system and the cDNA libraries were sequenced on an Illumina NextSeq instrument.

### Data processing and quality control

FASTQ sequence files were converted to Digital Count Conversion (DCC) format using Bruker’s GeoMx NGS Pipeline (v.2.3.3.10) performing read adapter trimming, alignment, and duplicate removal. All analyses were performed in R^23^ using Bioconductor^24^. ROIs and probes, were defined by GeoMx configuration files and filtered based on standard QC metrics, including sequencing depth, probe detection rate, and background signal. ROIs failing QC thresholds were excluded from downstream analysis.

Filtered expression data were Q3 normalized across all remaining ROIs. Differential gene expression was performed using the GeomxTools mixedModelDE function, thresholds were set at a False Discovery Rate (FDR) ≤ 0.05 and fold change (FC) ≥ 1.5. Gene Ontology (GO) over-enrichment analysis and Gene Set Enrichment Analysis (GSEA) was performed using the enrichGO and GSEA functions, respectively, from the clusterProfiler analysis library^25^.

### Single Nuclei RNA sequencing

Single nuclei suspensions were isolated from lung tissue using the 10X genomics nuclei isolation protocol, and processed through a Chromium X, following the GEM-X 5’ v3 chemistry 10x genomics protocol^26^. Sequencing reads were mapped to the human genome build hg38 using Cell Ranger v8.0.0 (10x Genomics)^27^ to generate barcode matrices. Data were normalized, checked for quality, and processed using Scanpy^28^. Batch correction was performed using the sample as covariate using single-cell variational inference^29^. Cell type annotations were assigned using canonical marker genes based on previously published single-cell lung atlases^15,16,20,30^. Cell type dotplots, Uniform Manifold Approximation and Projection (UMAP) plots, and cell counts were generated using Scanpy^28^. Differential gene expression for the single-nuclei data was performed using MAST^31^, differentially expressed gene (DEG) analysis thresholds were adjusted (FDR≤0.1; FC≥1.25) to account for differences in dataset-specific variability. ≥A 1.25 FC cutoff was chosen due to a minimal number of DEGs when using ≥1.5 FC (data not shown).

### Serum Proteomics

We leveraged a published serum proteomic dataset of 237 relatives of patients with IPF from two independent cohorts in which 26% had ILA^32^. The training cohort consisted of subjects from the GCS-PF cohort, a longitudinal study of first-degree relatives of patients with pulmonary fibrosis conducted at Brigham and Women’s Hospital (BWH) and BCM. In this cohort, 49 ILA and 107 Non-ILA subjects were identified through CT scans and included in the analysis. The protocols were approved by the IRBs of BWH (2016P000837) and BCM (H-46741). A validation cohort consisted of 13 ILA and 68 Non-ILA subjects from the University of Colorado and National Jewish Health^5^ (COMIRB #15–1147; NJH IRB 1441a). Plasma proteins were measured using the Somalogic SomaScan v4.1 proteomic platform as previously described^32^.

Using a log10 normalized and centered protein expression, we calculated differential protein expression between ILA and Non-ILA subjects. The list of differentially expressed proteins (p<0.05) was used to filter each compartment and cell type gene signature to proteins that were differentially expressed in plasma. These refined gene signatures were used in Least Absolute Shrinkage and Selection Operator (LASSO) based feature selection to derive a set of core proteins for each signature^33^. LASSO-regularized logistic regression models were used to derive reduced protein signatures.

## RESULTS

### Histologically Preserved Spatial Lung Niches in Preclinical FPF Demonstrate Compartment-Specific Transcriptional Abnormalities

To determine whether molecular abnormalities precede histologic remodeling in fibrosis, we performed spatial transcriptomic profiling of histologically preserved alveolar, airway, and vascular compartments from subjects with preclinical FPF and controls (**Figure 1A**). Representative ROIs confirmed preserved alveolar architecture, airway structures, and vascular compartments selected for analysis (**Figure 1B, 1E**, and **1H**).

**Figure 1.**
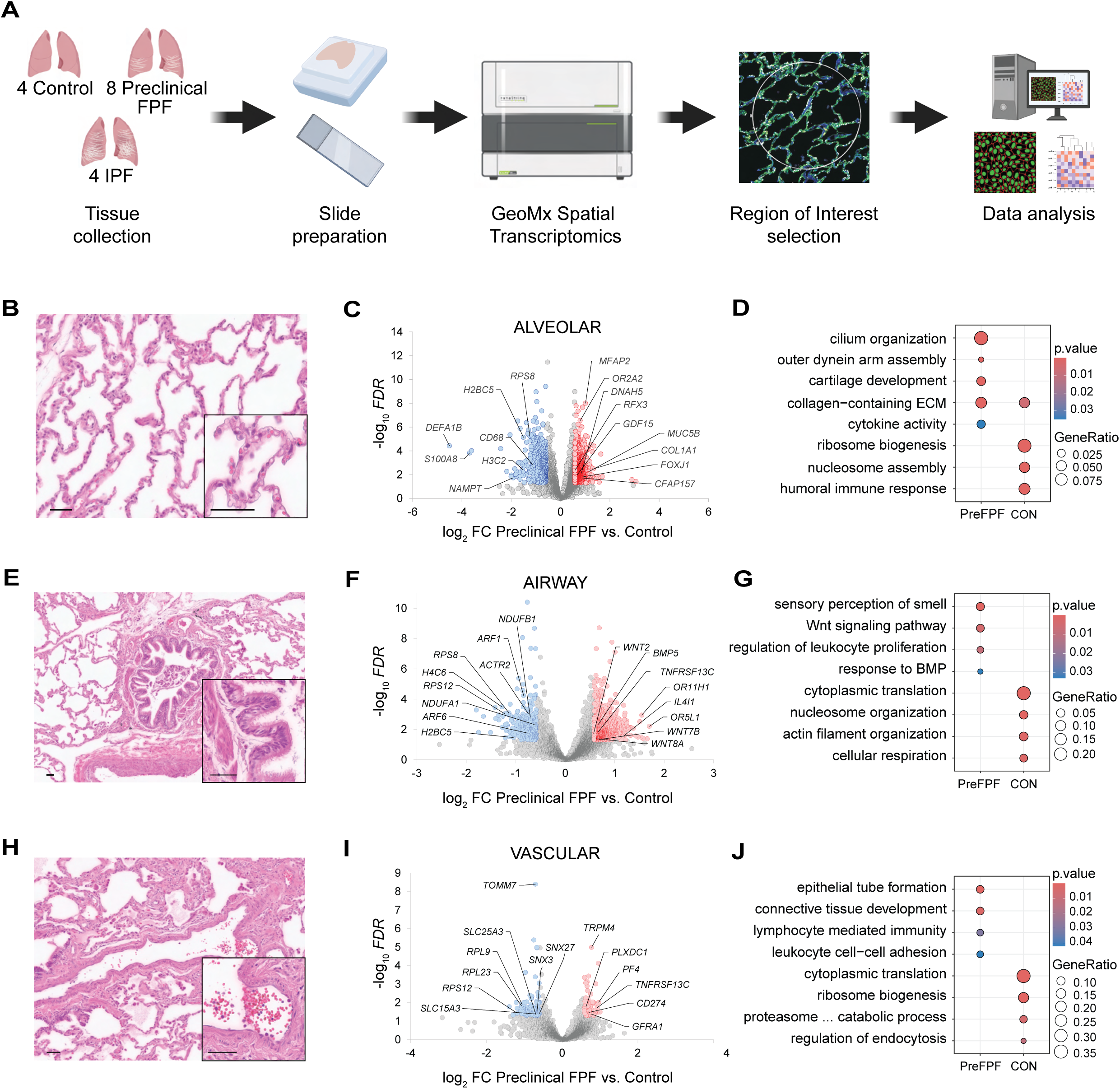
Histologically preserved lung compartments in preclinical FPF demonstrate compartment-specific transcriptional abnormalities. (A) Schematic overview of tissue acquisition, GeoMx spatial transcriptomic workflow, ROI selection, and downstream transcriptomic analyses. Histologically preserved alveolar, airway, and vascular compartments from subjects with preclinical FPF (Pre FPF) were compared with controls (CON). (B, E, H) Representative histologic images of preserved alveolar, airway (bronchioles), and vascular (arteries) compartments selected for spatial transcriptomic profiling. (C, F, I) Volcano plots of differentially expressed genes in preserved alveolar, airway, and vascular compartments from preclinical FPF compared with controls. Red dots (*right side*) represent higher expression in preclinical FPF; Blue dots (*left side*) represent higher expression in Control. (D, G, J) GO enrichment analyses of preserved alveolar, airway, and vascular compartments revealed both shared and compartment-specific pathway enrichment. ECM = Extracellular matrix **Figure 1.** Figure showing the spatial transcriptomic analysis of preserved alveolar, airway, and vascular compartments in preclinical FPF. Histologic images, differential gene expression, and pathway analyses identify molecular changes compared with control lungs.

Differential expression analyses identified substantial transcriptional abnormalities across preserved lung compartments from preclinical FPF subjects (**Figures 1C, 1F**, and **1I**). Despite overlap across compartments, distinct compartment-specific signatures were observed. Preserved alveolar regions displayed ECM-, epithelial-, and ciliary-associated programs (*i.e.,* COL1A1, MUC5B, MFAP2, FOXJ1, CFAP157, DNAH5). Preserved airways (bronchioles) demonstrated altered developmental (*i.e.,* WNT2, WNT7B, WNT8A, and BMP5) signaling pathways. Preserved arteries demonstrated altered vascular, connective tissue, and immune-associated pathways (*i.e.,* PF4, PLXDC1, CD274, and TNFRSF13C).

GO analyses revealed both shared and compartment-specific pathway enrichment across preserved lung compartments (**Figures 1D, 1G,** and **1J**). Enrichment was observed in alveolar regions for collagen-containing ECM and cilium organization; in airway regions for Wnt signaling, response to Bone Morphogenetic Protein, and actin filament organization; and in arterial regions for connective tissue development, leukocyte cell-cell adhesion, lymphocyte-mediated immunity, and epithelial tube formation pathways. These data reveal that histologically preserved alveolar, airway, and vascular compartments in preclinical FPF exhibit spatially organized compartment-specific molecular abnormalities.

### Remodeled Regions from Preclinical FPF and IPF Exhibit Overlapping and Distinct Molecular Programs

We next used spatial transcriptomics to compare fibroblastic foci, honeycombing regions, remodeled distal airways, and arteries with intimal hyperplasia from preclinical FPF and advanced IPF (**Figures 2A, 2D, 2G,** and **2J**).

**Figure 2.**
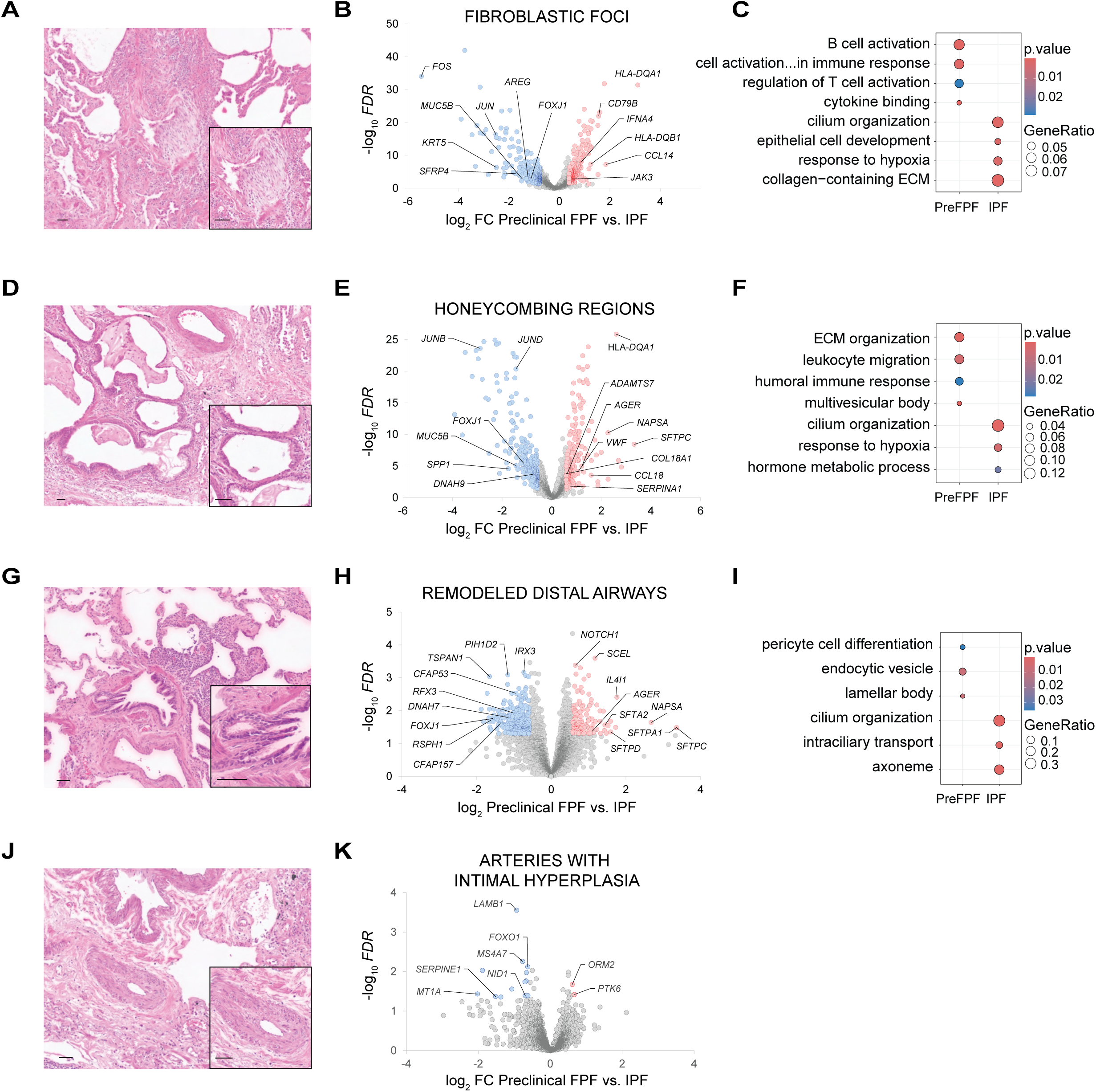
Remodeled regions from preclinical FPF and IPF exhibit overlapping and distinct molecular programs. (**A, D, G, J**) Representative histologic images of fibroblastic foci, honeycombing regions, remodeled distal airways, and arteries with intimal hyperplasia selected for GeoMx spatial transcriptomic profiling from subjects with preclinical FPF (Pre FPF) and IPF. (**B, E, H, K**) Volcano plots of differentially expressed genes between corresponding remodeled regions from preclinical FPF and advanced IPF. Red dots (*right side*) represent higher expression in preclinical FPF; Blue dots (*left side*) represent higher expression in IPF. (**C, F, I**) GO enrichment analyses demonstrated both shared and compartment-specific pathway enrichment across corresponding remodeled regions (fibroblastic foci, honeycombing, and remodeled airways) from preclinical FPF and IPF. Arteries with intimal hyperplasia did not yield sufficient differentially expressed genes to support GO enrichment analysis. **Figure 2.** Figure comparing remodeled lung regions in preclinical FPF and advanced IPF. Histologic images, differential gene expression, and pathway analyses identify shared and distinct molecular changes across disease stages.

Correlation analyses demonstrated strong concordance between remodeled regions from preclinical FPF and advanced IPF across fibroblastic foci, honeycombing regions, remodeled airways, and vascular compartments (**Figure S1**). Despite this overlap, differential expression analyses identified distinct transcriptional differences between remodeled regions of preclinical FPF vs. advanced IPF (**Figures 2B, 2E, 2H,** and **2K**), occurring primarily within fibroblastic foci, honeycombing regions, and remodeled airways. Fibroblastic foci from preclinical FPF were enriched for alveolar epithelial and surfactant-associated programs (*i.e.,* AGER, NAPSA, and SFTPC), whereas advanced IPF fibroblastic foci exhibited increased expression of ciliary- and ECM-associated transcripts (*i.e.,* MUC5B, FOXJ1, DNAH9, and SPP1).

Honeycombing regions from preclinical FPF exhibited differential expression of immune-associated and ECM-associated programs, while those from advanced IPF displayed reduction of alveolar epithelial and ciliary programs. Remodeled airways from preclinical FPF had increased expression of epithelial and developmental signaling-associated genes (*i.e.,* NOTCH1, AREG, MUC5B, SFRP4, and KRT5), whereas those from advanced IPF showed reduced alveolar epithelial-associated transcripts. Arteries with intimal hyperplasia showed only modest transcriptional differences between preclinical FPF vs. advanced IPF.

GO analyses revealed shared and compartment-specific pathway enrichment across remodeled regions from preclinical FPF and advanced IPF (**Figures 2C, 2F,** and **2I**). Pathway enrichment was observed in fibroblastic foci for epithelial development, ECM, and immune-associated pathways; in honeycombing regions for ECM organization, leukocyte migration, humoral immune response, and hypoxia-associated pathways; and in remodeled airways for cilium and epithelial-associated pathways. Thus, remodeled regions from preclinical FPF and advanced IPF share overlapping molecular programs while retaining compartment-specific transcriptional features.

### Paired Spatial Analyses in FPF Subjects Demonstrate Distinct Molecular Programs Between Preserved and Remodeled Regions

To define spatially organized molecular differences within preclinical FPF lungs, we performed paired spatial transcriptomic analyses comparing preserved and remodeled regions within the same tissue specimens. Preserved alveolar regions were compared with fibroblastic foci and honeycombing regions, while preserved distal airways were compared with remodeled distal airways.

Differential expression analyses identified distinct compartment-specific molecular programs between paired preserved and remodeled regions within preclinical FPF lungs (**Figures 3A, 3C,** and **3E**). Fibroblastic foci demonstrated increased ECM-associated and stromal-associated programs (*i.e.,* COL1A1, COL1A2, COL3A1, CTHRC1, SFRP2, MMP2), whereas paired preserved alveolar regions retained alveolar epithelial and surfactant-associated signatures (*i.e.,* AGER, SFTPC, ABCA3, NAPSA, CLDN18). Honeycombing regions exhibited increased expression of ECM-associated, hypoxia-associated, and stromal-associated genes (*i.e.,* SFRP2, SERPINF1, THBS1, LTBP1, and IGF1) relative to paired preserved alveolar regions, which retained endothelial- and alveolar epithelial-associated programs (*i.e.,* EPAS1, CAV1, CLIC5, and AGER).

**Figure 3.**
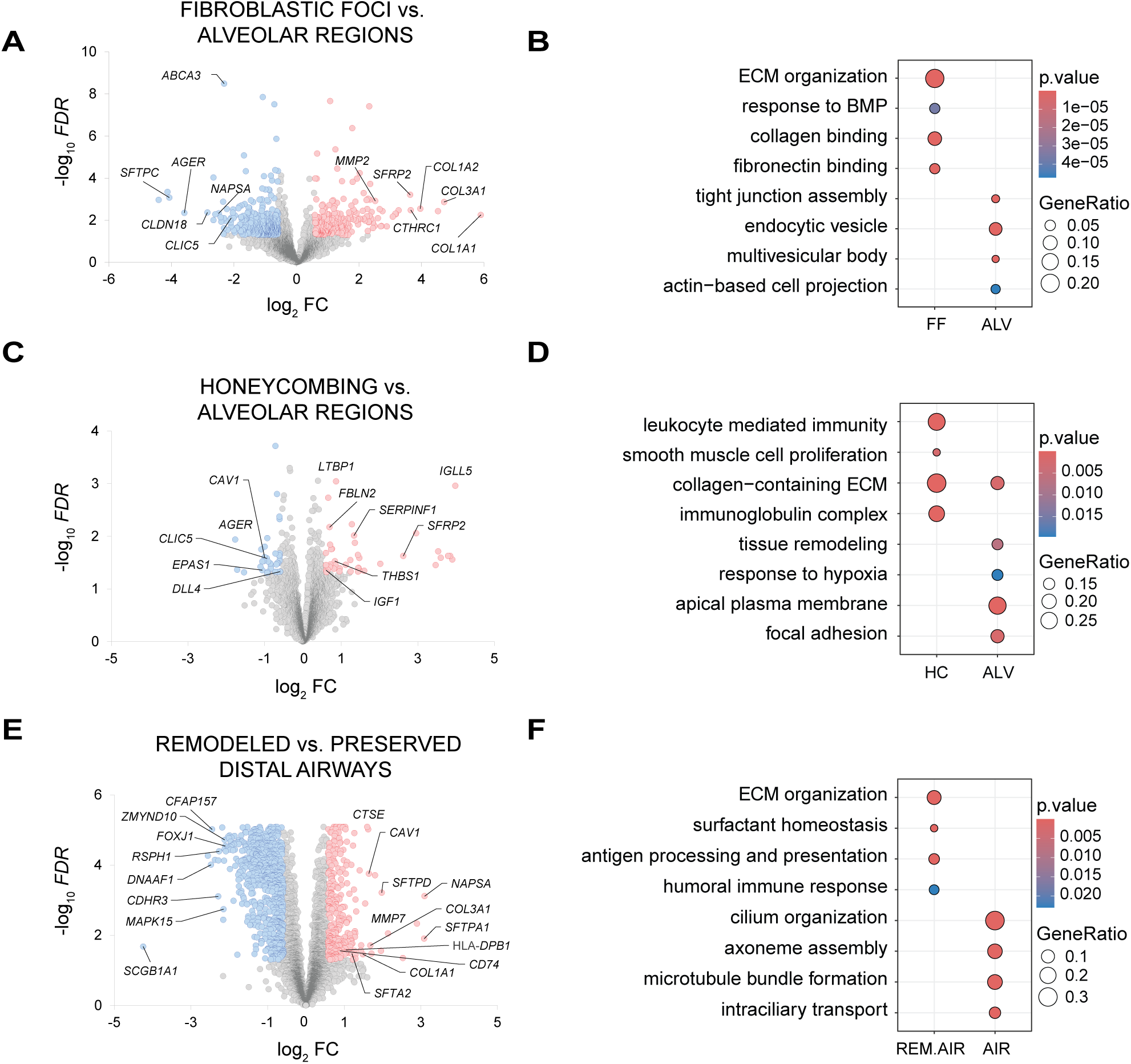
Paired spatial analyses identify compartment-specific molecular differences between preserved and remodeled regions in preclinical FPF. (**A, C, E**) Volcano plots of differential gene expression between paired preserved and remodeled regions within the same preclinical FPF (Pre FPF) lung specimens. Fibroblastic foci (FF) and honeycombing (HC) regions were compared with paired preserved alveolar (ALV) regions, while remodeled distal airways (REM.AIR) were compared with paired preseverved distal airways (AIR). Red dots (*right side*) represent higher expression in the remodeled region; Blue dots (*left side*) represent higher expression in the preserved region. (**B, D, F**) GO analyses of compartment-specific pathway enrichment across paired preserved and remodeled regions within preclinical FPF lungs (fibroblastic foci and honeycombing regions vs. paired preserved alveolar regions; remodeled vs. preserved distal airways). **Figure 3.** Figure showing paired comparisons of preserved and remodeled regions in preclinical FPF. Volcano plots identify differentially expressed genes, and gene ontology analyses reveal biological pathways associated with regional tissue remodeling.

Remodeled distal airways exhibited increased ECM-associated, epithelial-associated, and immune-associated programs (*i.e.,* COL1A1, COL3A1, MMP7, ICAM1, and VIM), whereas paired preserved distal airways retained ciliary-associated and epithelial homeostatic signatures (*i.e.,* FOXJ1, RSPH1, DNAAF1, CFAP157, CDHR3, and MAPK15).

GO analyses revealed compartment-specific pathway enrichment across paired preserved and remodeled regions within preclinical FPF lungs (**Figures 3B, 3D,** and **3F**). Enrichment was observed in fibroblastic foci for focal adhesion, smooth muscle cell proliferation, and hypoxia-associated pathways; in honeycombing regions for fibroblast proliferation and focal adhesion; and in remodeled distal airways for antigen processing and presentation, humoral immune response, and surfactant homeostasis. Paired preserved distal airways were enriched for cilium-related processes.

### Single-Nucleus RNA Sequencing Identifies Epithelial Cell States Associated with Preclinical FPF

To interrogate cellular programs underlying the compartment-specific molecular abnormalities identified by spatial transcriptomics, we performed snRNAseq in subjects with preclinical FPF included in the GeoMx analyses. These studies were performed in parallel with control and IPF samples to characterize epithelial cell states (see **Figure S2** for annotation) across the pulmonary fibrosis continuum (**Figure 4A**).

**Figure 4.**
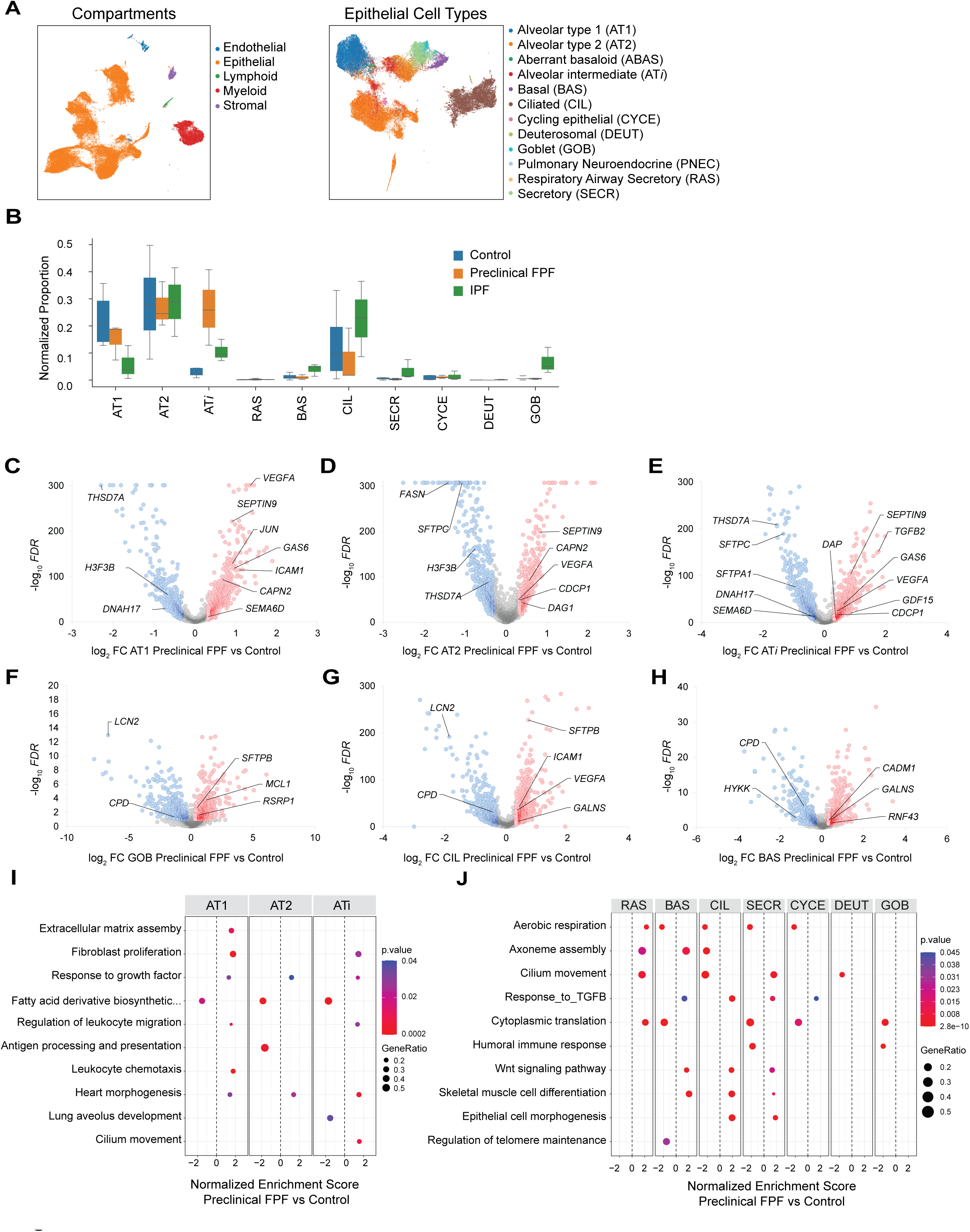
Single-nucleus RNA sequencing identifies epithelial cell states associated with preclinical familial pulmonary fibrosis. (**A**) UMAP visualization of epithelial cell populations identified by snRNAseq from control, preclinical FPF and IPF subjects. (**B**) Relative proportions of epithelial cell populations across control, preclinical FPF, and IPF lungs demonstrating altered epithelial cell-state composition. Differential expression analyses of distal alveolar epithelial populations including Alveolar epithelial type 1 (AT1) (**C**), alveolar epithelial type 2 (AT2) (**D**), and alveolar intermediate (AT*i*) (**E**) cells from preclinical FPF lungs compared with controls. Differential expression analyses of airway epithelial populations including goblet (GOB) (**F**), ciliated (CIL) (**G**), and basal (BAS) (**H**) cells from preclinical FPF lungs compared with controls, showing altered expression of epithelial, ciliary, and developmental signaling-associated genes. GO enrichment analyses of **(I)** distal alveolar and (**J**) airway epithelial populations. RAS = respiratory airway secretory; SECR = secretory; CYCE = cycling epithelial; DEUT = deuterosomal **Figure 4.** Figure showing single-nucleus RNA sequencing analysis of epithelial cell states in control, preclinical FPF, and IPF lungs. Cell population and differential expression analyses identify altered epithelial composition and molecular programs associated with early disease

Analysis of snRNAseq data revealed altered epithelial cell-state composition across preclinical FPF and IPF lungs (**Figure 4B**). AT1 epithelial populations demonstrated relative reduction across disease states. Preclinical FPF lungs demonstrated relative enrichment of alveolar intermediate (AT*i*) epithelial populations compared with controls. IPF lungs demonstrated increased representation of basal and ciliated epithelial populations. Differential expression analyses of distal alveolar epithelial populations identified ECM-associated, stress-response, epithelial transitional-associated, and growth factor signaling programs across AT1, AT2, and AT*i* populations (**Figures 4C, 4D, and 4E**). AT*i* cells demonstrated increased expression of epithelial transitional-associated and stress-response genes (*i.e.,* GDF15, KRT17, and GPX2). AT1 and AT2 populations demonstrated altered surfactant- and epithelial-associated homeostatic programs.

Proximal airway epithelial populations similarly demonstrated altered epithelial-, ciliary-, and developmental signaling-associated transcriptional programs in preclinical FPF lungs compared with controls (**Figures 4F, 4G, and 4H**). GO analyses revealed enrichment of ECM organization, fibroblast proliferation, leukocyte migration, epithelial differentiation, ciliogenesis, epithelial morphogenesis, WNT- and TGFβ-associated pathways across distal alveolar and airway epithelial populations (**Figures 4I and 4J**). Together, these findings support that preclinical FPF involves altered epithelial cell-state composition and enrichment of alveolar intermediate and airway epithelial programs associated with ECM-associated, stress-response, ciliary-associated, and developmental signaling pathways.

### Compartment and Cell-Defined Preclinical FPF Transcriptional Signatures are Partially Represented in the Peripheral Blood

To determine whether tissue-derived molecular programs identified in preclinical FPF are represented in the peripheral circulation, we repurposed a published peripheral blood proteomic cohort of first-degree relatives with and without ILAs and projected compartment-specific and epithelial transcriptional signatures into the circulating proteome.

Compartment-specific transcriptional signatures identified by spatial transcriptomics demonstrated limited overlap with circulating proteins and did not significantly associate with CT scan-based ILA status in the proteomic cohort (data not shown). To evaluate whether proteins shared between the tissue transcriptomic and circulating proteomic datasets improved discrimination, transcriptional signatures were refined to proteins also differentially expressed between ILA and Non-ILA subjects (**Figures 5A**). Models utilizing these overlapping protein sets distinguish ILA from Non-ILA subjects across compartments and epithelial cell types (**Figure 5A** and **5E**), except for the vascular compartment. For example, spatial transcriptomics identified 1138 DEG in the alveolar regions, of which 326 had corresponding proteins reflected in the peripheral blood and only 41 were differentially expressed between ILA and Non-ILA subjects in the protein samples.

**Figure 5.**
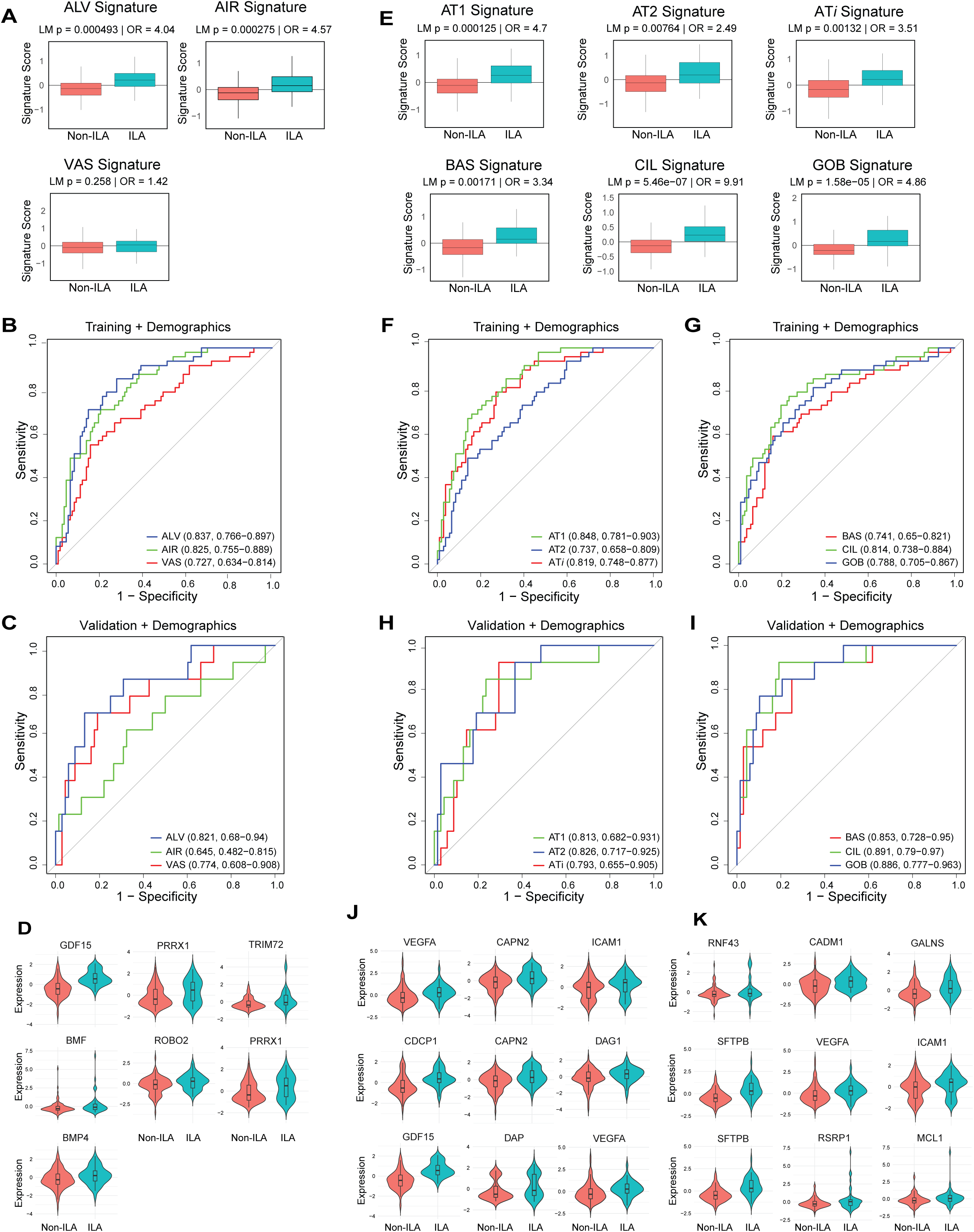
Spatially derived transcriptional signatures identify candidate circulating biomarkers in subjects with ILA. Lung-derived transcriptional signatures were projected into a previously published peripheral blood proteomic cohort of first-degree relatives of patients with IPF, with and without ILAs. (**A**) Boxplots displaying the mean compartment signature score in ILA or Non-ILA subjects. For each signature, a mean expression value score was calculated across all proteins within the signature and used to evaluate signature to ILA status association using linear regression models. Odds ratio and p values report the outcome of the association analysis. (**B-C**) ROC curves from post LASSO multivariable regression of a selected set of core proteins in the alveolar, airway and vascular compartments. Regression models were adjusted for demographics and batch effects in **(B)** Training and (**C**) Validation cohorts. (**D**) Violin plots of serum-detected proteins prioritized through LASSO analysis across histological compartments: Alveolar: GDF15, PRRX2 and TRIM72; Airway: BMF, ROBO2 and PRRX1; Vascular: BMP4 (**E**) Boxplots displaying the mean cell-based signature score in ILA or Non-ILA subjects. (**F-I**) ROC curves from post LASSO multivariable regression of a selected set of core proteins in epithelial cells from the proximal and distal regions. Regression models were adjusted for demographics and batch effects in (**F, G**) Training and (**H, I**) Validation cohorts which demonstrated consistent peripheral representation of selected tissue-informed proteomic signatures. (**J, K**) Violin plots of serum-detected proteins prioritized through LASSO analysis across epithelial cell-state signatures. (**J**) AT1: VEGFA, CAPN2, and ICAM1; AT2: CDCP1, CAPN2, and DAG1; AT*i*: GDF15, VEGFA, and DAP. (**K**) Basal: RNF43, CADM1, and GALNS; Ciliated: SFTPB, VEGFA, and ICAM1; Goblet: SFTPB, RSRP1, and MCL1. **Figure 5.** Figure showing associations between lung-derived transcriptional signatures and circulating blood proteins in individuals with and without ILA. Signature scores, predictive models, and prioritized proteins identify tissue-based candidate blood biomarkers of early pulmonary fibrosis.

To determine feature selection, we used LASSO, through which a prioritized set of core overlapping proteins was identified and evaluated using linear modeling. Proteomic signatures derived from alveolar and airway compartments demonstrated the strongest discrimination between subjects with ILAs and unaffected relatives, while vascular-associated signatures demonstrated comparatively modest peripheral representation (**Figure 5B**). These findings suggest that epithelial programs contribute significantly to circulating molecular abnormalities in preclinical pulmonary fibrosis. External validation in an independent cohort demonstrated consistent performance across compartments (**Figure 5C**). Serum-detected proteins identified through LASSO analysis demonstrated distinct compartment-specific profiles, with alveolar regions enriched for GDF15, PRRX1, and TRIM72; airway regions for BMF, ROBO2, and PRRX1; and vascular regions for BMP4 (**Figure 5D**).

Integration of snRNAseq data with circulating proteomics suggests that epithelial cell-state programs identified in preclinical FPF lungs were only partially represented in the peripheral blood. However, refined tissue-based protein sets identified in ILA lungs remained detectable in the circulation (**Figure 5E**). AT1 and AT*i* epithelial-associated programs demonstrated significant peripheral representation in subjects with ILAs. Linear models derived from the refined overlapping tissue-based proteome allowed for discrimination of ILA status across multiple epithelial cells in the training cohort, including AT1 (AUC 0.848, 95% CI 0.786–0.904), AT*i* (AUC 0.819, 95% CI 0.747– 0.880), and ciliated cells (AUC 0.814, 95% CI 0.736–0.884) (**Figures 5F** and **5G**).

External validation in an independent cohort demonstrated consistent performance across multiple epithelial cells, including ciliated, basal, and AT2 signatures (**Figures 5H** and **5I**). Serum-detected proteins prioritized through LASSO analysis demonstrated distinct epithelial cell-state protein profiles (**Figures 5J** and **5K**). AT1 cell signatures included VEGFA, CAPN2, and ICAM1, consistent with epithelial injury signaling. AT2 cell signatures prioritized CDCP1, CAPN2, and DAG1, reflecting epithelial stress and altered epithelial–matrix interactions.

AT*i* signatures prioritized GDF15, VEGFA, and DAP, supporting transitional injury states. Basal cell signatures prioritized RNF43, CADM1, and GALNS, consistent with epithelial remodeling. Ciliated cell signatures included SFTPB, VEGFA, and ICAM1, reflecting epithelial dysfunction and inflammatory signaling. Goblet cell signatures included SFTPB, RSRP1, and MCL1, suggesting aberrant epithelial differentiation. Together, these findings identify spatially and cell-defined molecular programs in preclinical FPF lungs, which are partially represented in the peripheral circulation.

## DISCUSSION

Here, we performed integrated spatial transcriptomic, single-cell, and peripheral blood proteomic profiling to define compartment- and cell-specific molecular programs associated with preclinical FPF. We demonstrate that early pulmonary fibrosis is characterized by biologically active molecular alterations across alveolar, airway, and vascular niches that precede fibrotic tissue remodeling. These observations extend prior radiographic and biomarker studies of ILAs by demonstrating that early fibrotic lung disease is defined not only by subtle imaging abnormalities, but also by coordinated molecular programs localized to discrete histologic compartments. Although ILAs identify individuals at increased risk for progression, radiographic phenotyping does not define the underlying cellular programs or compartment-specific pathways that ultimately influence disease trajectory and therapeutic responsiveness. The marked variability in compartment- and cell-specific molecular programs observed across preclinical FPF lungs suggests that biologic heterogeneity emerges early in a spatially organized disease process^34^.

Additionally, we identified epithelial-associated molecular programs within histologically preserved lung regions. Integration of spatial and snRNAseq profiling demonstrated enrichment of alveolar and airway epithelial states associated with ECM-related pathways and transitional epithelial phenotypes. Thus, epithelial dysfunction and aberrant repair responses arise early in pulmonary fibrosis before extensive fibrotic tissue remodeling^21^. Notably, epithelial-associated molecular programs localized to discrete alveolar and airway niches, suggesting that early fibrotic remodeling may emerge through spatially organized epithelial injury responses rather than diffuse homogeneous tissue injury and repair. These findings highlight the central role of epithelial dysregulation in early fibrotic lung disease.

Our findings inform the interpretation of circulating biomarkers associated with early pulmonary fibrosis. Circulating proteins have been associated with ILAs and progression of fibrotic lung disease, including matrix metalloproteinases, surfactant-associated proteins, and epithelial stress-response mediators, however, their relationship to compartment-specific lung biology remains unclear. Here, tissue-derived molecular programs identified within alveolar, airway, and vascular compartments demonstrated modest peripheral representation. Spatial transcriptomic profiling interrogated ∼12,000 genes across tissue compartments, whereas the SOMAscan platform measured ∼7,500 circulating proteins, representing a fraction of the circulating proteome. Many tissue-associated transcripts encode intracellular, structural, or ECM proteins that are not secreted or detectable in the circulation.

Furthermore, the peripheral blood proteome reflects contributions from multiple organs and vascular cell populations, which likely dilutes localized lung-derived molecular signals. Nevertheless, several epithelial-associated molecular programs demonstrated reproducible peripheral representation and were associated with ILA status. These findings suggest that circulating biomarkers may partially reflect biologically active tissue niches and epithelial cell states. However, the current data does not support definitive peripheral blood molecular endotyping of early pulmonary fibrosis. Instead, these observations provide evidence supporting the feasibility of tissue-informed biomarker development.

There are several limitations of the current study. First, lung tissue and peripheral blood analyses were performed in independent cohorts, limiting direct correlation between tissue-derived molecular programs and circulating biomarkers. Future paired lung-blood multi-omic studies using larger prospective cohorts and expanded proteomic platforms will determine whether circulating biomarkers can reproducibly represent tissue-defined molecular programs and ultimately support biologically meaningful endotyping of early pulmonary fibrosis. Second, the limited availability of preclinical lung tissue prevented definitive identification of molecular subtypes associated with disease progression. Third, although spatial transcriptomic profiling identified molecular programs across compartments, the single-nuclei analysis was enriched for epithelial populations, limiting comprehensive characterization of mesenchymal, endothelial, and immune cell states. Finally, the targeted nature of the spatial and single-nuclei platforms limited interrogation of multicellular interactions within discrete histologic niches. Larger prospective studies integrating spatial transcriptomic, proteomic, genomic, and longitudinal clinical data will determine how tissue-specific molecular programs influence clinical outcomes and response to therapy.

## Conclusions

Early pulmonary fibrosis is characterized by biologically active niche- and cell-specific molecular programs that are detectable before the development of established fibrotic tissue remodeling. Spatial transcriptomic profiling identified distinct molecular alterations across alveolar, airway, and vascular compartments, while single-cell analysis demonstrated prominent epithelial cell states associated with early fibrotic biology. Although tissue-derived molecular programs exhibited only modest peripheral representation, several epithelial-associated signatures were reproducibly associated with ILA status, supporting the feasibility of tissue-informed biomarker development. Our findings support biologically informed investigation of early pulmonary fibrosis and will facilitate future prospective studies integrating paired tissue and peripheral blood multi-omic profiling, that will define clinically meaningful molecular endotypes and enable precision therapeutic approaches.

## Author contributions

Study design: A.W. and I.O.R. Data acquisition, analysis, or interpretation: A.W., S.A.O., J.A.V., J.R., J.D.C.C., J.D.Z., N.J.M., M.E.R.E., C.H. Critical revision of the manuscript for important intellectual content: C.H., L.J.C., K.T., L.F.G.C., A.G.C., S.W.R., W.J.I., N.K., D.A.S., B.A.R., G.M.H., B.R.G., A.J. Bioinformatic analysis: S.A.O., M.E.R.E, C.C.

## Ethical approval/Patient consent statement

Protocols and consent forms for the collection of lung samples were approved by the Institutional Review Board (IRB) of the National Heart, Lung and Blood Institute and the National Human Genome Research Institute of the National Institutes of Health in Bethesda, Maryland (Protocols 99-H-0068, 04-HG-0211 and/or NCT00084305) or Baylor College of Medicine (BCM) (IRB #H-46823). Protocols for the collection of peripheral blood samples were approved by the IRBs of Brigham and Women’s Hospital (BWH) (2016P000837), BCM (H-46741), University of Colorado (COMIRB #15–1147) and National Jewish Health (NJH IRB 1441a). All subjects gave written consent as per the Declaration of Helsinki.

## Acknowledgements

We thank the patients and families who participated in our research protocols for their invaluable contributions to this work. Elements of the results from this study were presented at the American Thoracic Society International Conference in May 2025 in San Francisco.

