## Supplemental methods, tables and figures for "Early Pulmonary Fibrosis is Defined by Niche- and Cell-Specific Molecular Programs"

#### SUPPLEMENTARY METHODS

##### Lung tissue samples

Lung tissue was obtained from open lung biopsies of eight individuals with preclinical FPF, defined by radiographic evidence of ILA and absence of clinical symptoms. These were collected as part of a larger cohort of kindreds of patients with IPF[1] enrolled in protocols 99-H-0068, 04-HG-0211 and/or NCT00084305 of the National Heart, Lung, and Blood Institute and/or the National Human Genome Research Institute Institutional Review Boards. Their average age was 55.35 years, four were female and four were male. In addition, whole lung explants were collected from four individuals with end-stage IPF undergoing lung transplantation. The diagnosis of IPF was established according to the American Thoracic Society/European Respiratory Society (ATS/ERS) guidelines[2]. Control samples were obtained from organ donors whose lungs were rejected for transplantation to living recipients. IPF and Control lung samples were collected after approval by the Baylor College of Medicine Institutional Review Board (H-46823). Clinical and demographic characteristics of study subjects are listed in **Table S1.**

##### Bruker Spatial Biology GeoMx Digital Spatial Profiler

Lung sections were fixed by immersion in 10% neutral buffered formalin solution and embedded in paraffin. Thin (5 μm) sections were deparaffinized and rehydrated. Tissue slides were stained with hematoxylin-eosin (H&E) to evaluate histopathologic changes and facilitate selection of Regions of Interest (ROI). Spatial transcriptomic profiling was performed using a Bruker Spatial Biology GeoMx Digital Spatial Profiler (DSP). RNA slides were prepared according to the manufacturer's protocols and instructions. 5 μm tissue sections were hybridized with the GeoMx Whole Transcriptome Atlas probe set and stained with fluorescent antibodies (pan-cytokeratin, SYTO13 and CD45) provided within the GeoMx reagent kits. Slides were placed into the instrument and scanned at high resolution. ROI were selected by a board-certified and fellowship trained pulmonary pathologist (J.A.V.) guided by H&E staining and fluorescent antibody staining. A brief description of each ROI type is presented in **Table S2**. Serial ultraviolet illumination of each region was used to collect the probe barcodes into a ninety-six well collection plate. Each ROI was then indexed using an Illumina dual indexing system and the cDNA libraries were sequenced on an Illumina instrument.

Data processing and quality control

FASTQ sequence files were converted to Digital Count Conversion (DCC) format using Bruker's GeoMx NGS Pipeline (v.2.3.3.10) performing read adapter trimming, alignment, and duplicate removal. All analyses were performed in R[3] using Bioconductor[4]. Using the GeomxTools R package[5], ROIs and probes, as defined by the GeoMx configuration files Hs_R_NGS_WTA_v1.0.pkc, were filtered based on standard QC metrics, including sequencing depth, probe detection rate, and background signal. ROIs failing QC thresholds were excluded from downstream analysis.

Filtered expression data were Q3 normalized across all remaining ROIs. Differential gene expression was performed using a liner mixed-effect model to account for fixed effects (sample groups of interest) and random effects (GeoMx slide) using the

GeomxTools mixedModelDE function. Gene Ontology over-enrichment analysis and GSEA was performed using the enrichGO and GSEA functions, respectively, from the clusterProfiler analysis library[6].

**Single Nuclei RNA sequencing**

Single nuclei suspensions were generated from snap frozen tissue lysates, following the 10X genomics nuclei isolation protocol, and processed through a Chromium X following the GEM-X 5' v3 chemistry 10x genomics protocol[7]. Library quality control was assessed on an Agilent Bioanalyzer High Sensitivity DNA chip. cDNA libraries were sequenced on an Illumina platform. Sequencing reads were mapped to the human genome build hg38 using Cell Ranger v8.0.0 (10x Genomics)[8] to generate barcode matrices. Data was normalized, checked for quality, and processed using Scanpy[9]. Batch correction was performed using the sample as covariate using SCVI[10]. Cell type annotations were assigned using canonical marker genes based on previously published single-cell lung atlases[11–13]. Cell type dotplots (**Figure S2**), Uniform Manifold Approximation and Projection (UMAP) plots, and cell counts were generated using Scanpy[9.]. Differential expression analysis was performed across cell types and study conditions using a threshold of absolute fold change > 1.25 and FDR < 0.1 to define differentially expressed transcriptional signatures associated with each cell type.

**Serum Proteomics**

<u>Differential Protein Expression and Gene Signature Filtering:</u>

We leveraged a previously published serum proteomic dataset of 237 relatives of patients with IPF from two independent cohorts out of which 26% had ILA[14]. The training cohort was composed of subjects from the GCS-PF cohort, a longitudinal study of first-degree relatives of patients with pulmonary fibrosis conducted at Brigham and Women's Hospital (BWH) and Baylor College of Medicine (BCM). In this cohort, 49 subjects with ILA and 107 without ILA were identified through prone computed tomography scans to assess for the presence of ILAs and included in the analysis. The protocols for this cohort and the collection of samples were approved by the Institutional Review Board of BWH (2016P000837) and BCM (H-46741). A validation cohort consisted of 13 subjects with ILA and 68 without ILA identified as part of a cohort from the University of Colorado and National Jewish Health collected through a previously published protocol[15] approved by the Institutional Review Board (COMIRB #15–1147; NJH IRB 1441a).

Plasma proteins were measured through the Somalogic SomaScan v4.1 proteomic platform as previously described[14]. Using log10 normalized and centered protein expression data, we calculated differential protein expression between ILA and Non-ILA subjects. Briefly, using the R linear analysis package, limma[16], a linear model was fit to a group means parameterization design matrix which included adjustments for demographics (age, gender, and smoking status). Proteins considered differentially expressed had a $p < 0.05$. Using this list of differentially expressed proteins, each compartment and cell type gene signature was filtered to gene symbols also differentially expressed in plasma.

<u>LASSO feature selection:</u>

Proteomic data were organized into a sample-by-protein expression matrix and aligned to clinical metadata by sample identifier. No additional normalization or scaling was performed prior to analysis, as input data were previously log-transformed and standardized. Interstitial lung abnormality (ILA) status was encoded as a binary outcome variable. The refined GeoMx and cell type signatures were used to construct model-specific feature sets. For each signature, a predictor matrix was generated consisting of the corresponding proteins, with inclusion of demographic covariates (age, gender, and smoking history) encoded using model matrix expansion for categorical variables. Feature selection was performed using penalized logistic regression with LASSO regularization using the R package glmnet[17,18].

Models were trained using k-fold cross-validation, in which samples were partitioned into training and validation subsets. Within each fold, the LASSO model was fit on the training data with tuning of the regularization parameter (λ) via internal cross-validation. The optimal λ (λ_min) was selected, and predictions were generated for the held-out samples. Cross-validation was repeated across folds to obtain out-of-sample predicted probabilities for each sample. Model performance was evaluated using the area under the receiver operating characteristic curve (AUC), calculated from cross-validated predictions. Confidence intervals (95%) for AUC were estimated using nonparametric bootstrap resampling with 1,000 iterations, and percentile-based intervals were reported. Receiver operating characteristic (ROC) curves were generated for each model, and performance metrics were summarized in tabular form to facilitate comparison across signatures.

For validation, trained models were applied to an independent validation cohort. Model inputs were constructed using the same feature definitions and covariate encoding as in the training data. Predictions were generated without retraining, and performance was evaluated using ROC analysis and bootstrap-derived confidence intervals as described above.

Core protein evaluation:

Core protein signatures were derived from prior LASSO-based feature selection and evaluated as fixed multivariable models. For each predefined protein signature, a model matrix was constructed by selecting the corresponding analytes and augmenting them with demographic covariates (age, gender, and smoking history), encoded using a design matrix framework. No additional scaling or transformation was performed. Model performance in the training cohort was assessed using repeated k-fold cross-validation. In each fold, logistic regression models were fit using the training subset, and predictions were generated for the held-out samples.

Predicted probabilities of ILA were aggregated across all folds to produce out-of-sample predictions for each model. Discriminatory performance was evaluated using the area under the receiver operating characteristic curve (AUC). Confidence intervals for AUC were estimated via nonparametric bootstrap resampling, with percentile-based 95% confidence intervals derived from 1,000 resampling iterations. Receiver operating characteristic (ROC) curves were generated for each model, and AUC values with confidence intervals were displayed. Performance metrics were summarized in tabular form to enable comparison across protein signatures. For external validation, the same fixed protein signatures were applied to an independent validation cohort. Models were re-fit on the full training dataset and used to generate predicted probabilities in the validation cohort without modification. ROC analysis and bootstrap-based confidence intervals were computed as described above to assess generalizability.

#### SUPPLEMENTARY TABLE LEGENDS

**Table S1. Demographic and clinical characteristics of study subjects.**

Demographic and clinical characteristics of subjects included in the GeoMx spatial transcriptomics and single-nucleus RNA sequencing cohorts. Data are presented as mean ± Standard deviation or n (%). Pulmonary function test values are reported as percent predicted. Control samples were obtained from donor lungs. UIP: usual interstitial pneumonia; DLCO: diffusing capacity of the lung for carbon monoxide; FVC: forced vital capacity; FEV1: forced expiratory volume in one second.

**Table S2. Region of interest (ROI) selection strategy for spatial transcriptomic profiling.**

Description of selected ROIs and predefined spatial comparisons across control, ILA, and IPF lungs. ROIs were selected based on histopathologic features to capture preserved and remodeled anatomical niches associated with early and advanced fibrotic remodeling. These predefined spatial comparisons were used to investigate compartment-specific transcriptional programs across airway, alveolar, and vascular regions during fibrosis progression.

| Characteristic | Spatial Transcriptomics | | | Single nuclei RNA sequencing | | |
|---|---|---|---|---|---|---|
| | Control (n=4) | Preclinical FPF (n=8) | IPF (n=4) | Control (n=4) | Preclinical FPF (n=3) | IPF (n=3) |
| | n (%) | n (%) | n (%) | n (%) | n (%) | n (%) |
| **Age (mean ± SD)** | 50.3 ± 7.1 | 55.4 ± 8.4 | 62.8 ± 8.2 | 51.5 ± 7.1 | 58.7 ± 10.2 | 66.0 ± 6.1 |
| **Gender** | | | | | | |
| Male | 1 (25%) | 4 (50%) | 2 (50%) | 1 (25%) | 3 (100%) | 1 (33.3%) |
| Female | 3 (75%) | 4 (50%) | 2 (50%) | 3 (75%) | | 2 (66.7%) |
| **Race** | | | | | | |
| White | 2 (50%) | 7 (87.5%) | 2 (50%) | 3 (75%) | 2 (66.7%) | 2 (66.7%) |
| Black or African American | 1 (25%) | | 1 (25%) | 1 (25%) | | 1 (33.3%) |
| Asian | 1 (25%) | | | | | |
| Hispanic | | 1 (12.5%) | 1 (25%) | 1 (25%) | 1 (33.3%) | |
| **Ethnicity** | | | | | | |
| Hispanic or Latino | | 1 (12.5%) | 2 (50%) | 1 (25%) | 1 (12.5%) | 1 (33.3%) |
| Not Hispanic or Latino | 4 (100%) | 7 (87.5%) | 2 (50%) | 3 (75%) | 7 (87.5%) | 2 (66.7%) |
| **History of smoking** | | | | | | |
| Yes | 1 (25%) | NA | 2 (50%) | 2 (50%) | NA | 2 (66.7%) |
| No | 3 (75%) | NA | 2 (50%) | 2 (50%) | NA | 1 (33.3%) |
| **Antifibrotic therapy** | | | | | | |
| Nintedanib | | | 1 (25%) | | | 1 (33.3%) |
| Pirfenidone | | | 2 (50%) | | | 2 (66.7%) |
| **Pulmonary function tests (mean ± SD)** | | | | | | |
| FVC% | NA | 90.7 ± 14.1 | 50.2 ± 21.0 | NA | 95.2 ± 15.9 | 56.2 ± 22.0 |
| FEV1% | NA | 97.8 ± 14.4 | 54.2 ± 25.0 | NA | 104.9 ± 13.5 | 61.6 ± 26.3 |
| DLCO% | NA | 83.3 ± 20.2 | 32.5 ± 18.9 | NA | 72.6 ± 14.8 | 24.0 ± 9.5 |
| **Pathology** | | | | | | |
| Usual Interstitial Pneumonia (UIP) | | 5 (62.5%) | 4 (100%) | | 2 (66.7%) | 3 (100%) |
| Mild fibrosis | | 1 (12.5%) | | | | |
| Diffuse interstitial fibrosis | | 1 (12.5%) | | | 1 (33.3%) | |
| Bronchiolocentric interstitial pneumonia | | 1 (12.5%) | | | | |
| **Outcome** | | | | | | |
| Donor | 4 (100%) | | | 4 (100%) | | |
| Alive | | 1 (12.5%) | | | | |
| Death | | 3 (37.5%) | | | 2 (66.7%) | |
| Lung transplant | | 1 (12.5%) | 4 (100%) | | | 3 (100%) |
| Unknown | | 3 (37.5%) | | | 1 (33.3%) | |

**Table S1**

| Compartment | Control | Preclinical FPF or IPF |
|---|---|---|
| **Airway** | Non-cartilaginous distal airway (bronchiole) | Unremarkable non-cartilaginous distal airway (bronchiole) |
| | | Remodeled non-cartilaginous airway (bronchiole ), with adjacent bronchiolized parenchyma (bronchiolar metaplasia) |
| **Alveolar** | Alveolar regions | Histologically preserved alveolated parenchyma (unremarkable alveolar regions) |
| | | Honeycombing regions |
| | | Fibroblastic foci |
| **Vascular** | Small pulmonary arteries | Histologically preserved medium to small sized arteries |
| | | Medium-to-small sized arteries with intimal hyperplasia in fibrotic areas |

#### Table S2

#### SUPPLEMENTARY FIGURE LEGENDS

**Figure S1. Early-stage familial pulmonary fibrosis and late-stage IPF share transcriptional programs across multiple spatial regions.**

Correlation plots showing overlap of differential gene expression between ILA versus control and IPF versus control across fibroblastic foci, honeycombing regions, remodeled airways, and arteries with intimal hyperplasia. Differentially expressed genes demonstrated significant directional concordance between early-stage familial pulmonary fibrosis and late-stage IPF across multiple remodeled anatomical niches, supporting shared transcriptional programs associated with fibrotic remodeling when compared to controls.

**Figure S2. Canonical marker-based annotation of epithelial cell populations in the snRNAseq assay.**

(**A**) Dotplot showing epithelial cell type annotation using canonical marker genes. Dot size represents the proportion of cells expressing each gene, while color intensity reflects relative expression levels. Expression patterns across epithelial populations were used to identify major epithelial cell subtypes and transitional epithelial cell states supporting accurate classification of epithelial cell populations for downstream analysis.

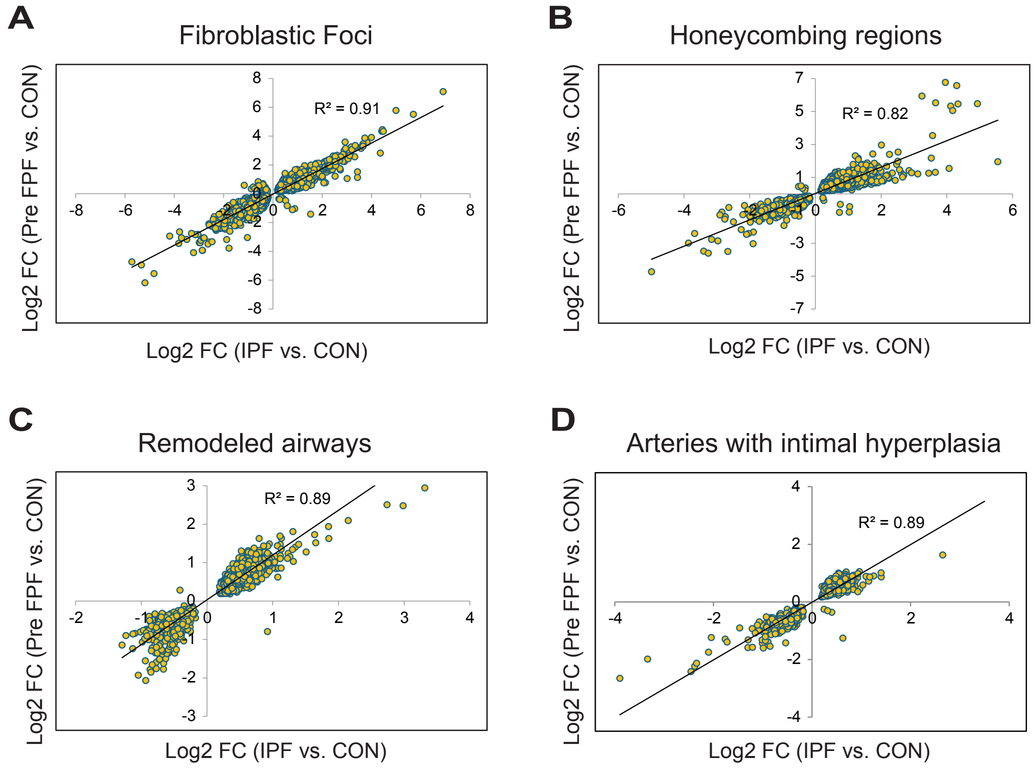
A
Fibroblastic Foci
R² = 0.91
Log2 FC (Pre FPF vs. CON)
Log2 FC (IPF vs. CON)
B
Honeycombing regions
R² = 0.82
Log2 FC (Pre FPF vs. CON)
Log2 FC (IPF vs. CON)
C
Remodeled airways
R² = 0.89
Log2 FC (Pre FPF vs. CON)
Log2 FC (IPF vs. CON)
D
Arteries with intimal hyperplasia
R² = 0.89
Log2 FC (Pre FPF vs. CON)
Log2 FC (IPF vs. CON)

**Figure S1**

A

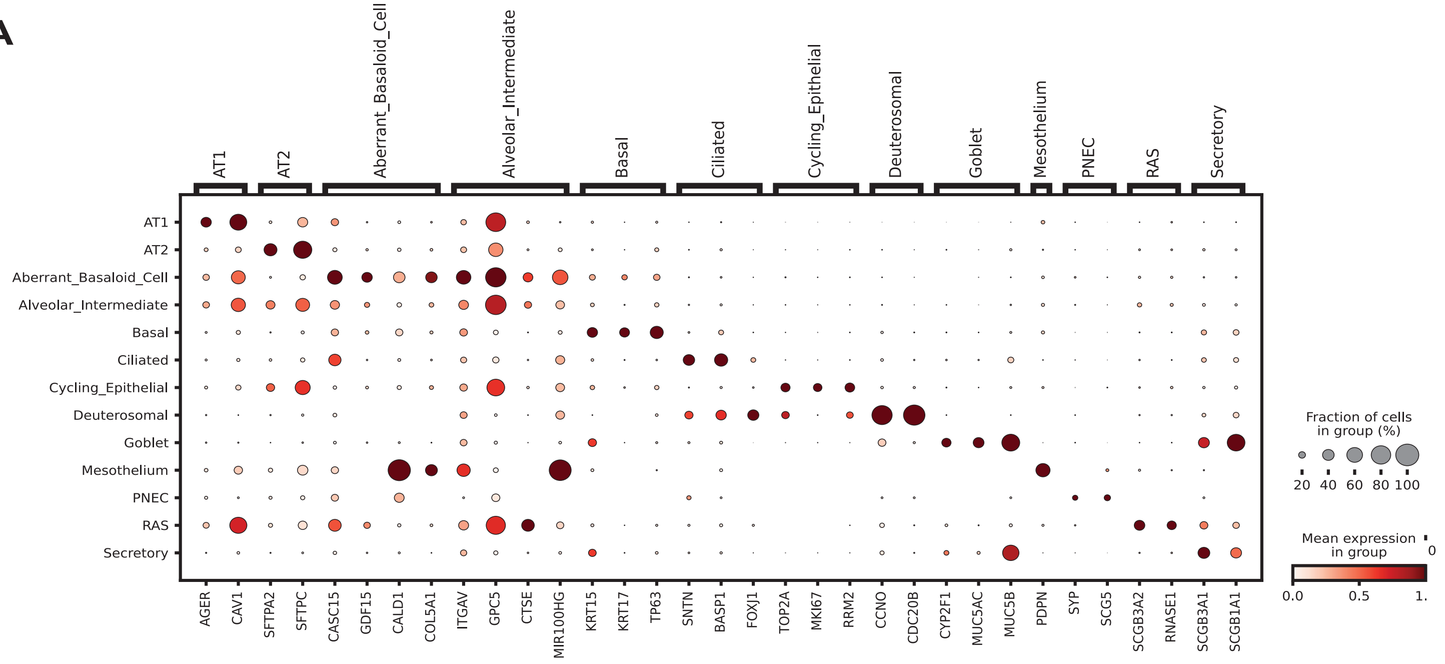

AT1
AT2
Aberrant_Basaloid_Cell
Alveolar_Intermediate
Basal
Ciliated
Cycling_Epithelial
Deuterosomal
Goblet
Mesothelium
PNEC
RAS
Secretory
AGER
CAV1
SFTPA2
SFTPC
CASC15
GDF15
CALD1
COL5A1
ITGAV
GPC5
CTSE
MIR100HG
KRT15
KRT17
TP63
SNTN
BASP1
FOXJ1
TOP2A
MKI67
RRM2
CCNO
CDC20B
CYP2F1
MUC5AC
MUC5B
PDPN
SYP
SCG5
SCGB3A2
RNASE1
SCGB3A1
SCGB1A1
Fraction of cells in group (%)
20 40 60 80 100
Mean expression in group
0
0.0 0.5 1.


### Figure S2
